# Linking somatic mutations in cancer to the electronic properties of DNA

**DOI:** 10.64898/2026.01.02.697387

**Authors:** Benoît de Witte, Cyril Karamaoun, Pauline Hermans, Maxime Tarabichi, Fabrizio Pucci, Marianne Rooman

## Abstract

Oxidative stress, generated by both endogenous and exogenous agents, can cause DNA lesions that, if not repaired, accumulate as somatic mutations and can contribute to cancer initiation. Here, we explored this problem through the lens of DNA electronic properties, quantified by the vertical ionization potential (vIP) of nucleobase motifs, which reflects their susceptibility to oxidation. We analyzed genome-wide experimental data on oxidative DNA damage and found that the highest damage levels occur in regions with low vIP values, supporting a causal link between them. The analysis of cancer mutational signatures and their annotated aetiologies revealed strong anticorrelations between mutation frequency and vIP values, particularly in cancers driven by oxidative DNA damage, such as lung cancer. We further computed anticorrelations between vIP values and the frequencies of mutated motifs across coding and non-coding regions and across different mutation types, observing the strongest anticorrelations for silent mutations, consistent with their reduced selective pressure. Moreover, similar anticorrelations were observed for somatic mutations in cancer and normal tissues, as well as for germline mutations, suggesting that they arise from similar mutagenesis processes. This work clarifies how oxidative damage, DNA electronic properties and carcinogenesis are related and help identify genomic regions more prone to mutations.

## 1 Introduction

Cancer initiation and progression are driven by cellular alterations primarily resulting from the progressive accumulation of somatic mutations within the genomes of normal tissues throughout an individual’s lifetime. These genetic changes disrupt normal cellular functions, ultimately leading to uncontrolled cell growth and division, and to tumor development.

But only some, called driver mutations, provide a selective advantage to the cells in which they occur, often by impairing the function of some key proteins that, for example, control the cell cycle or apoptosis [1]. These events facilitate the transition from a benign to a malignant lesion and promote genome instability. In contrast, passenger mutations also accumulate but do not confer any selective advantage to their host cells [2] and can in some cases even have slightly deleterious effects [3, 4].

Among the various types of mutations, single nucleotide substitutions (SBSs), in which one nucleobase is replaced by another, play a major role, and we will focus on them in this work. SBSs are primarily caused by the exposure of cells to a wide range of endogenous and exogenous agents, and lie at the heart of carcinogenesis [5]. SBSs are often caused by both exogenous exposures and endogenous agents, and lie at the heart of carcinogenesis [5]. There are multiple mechanisms of SBS formation, which depend on a wide variety of mutagenesis pathways. One example is the oxidation of DNA bases by reactive oxygen species (ROS), which are generated as byproducts of cellular metabolism [6, 7, 8]. Guanine is oxidized to 8-oxo-7,8-dihydroguanine (8-oxoG) at a high rate in the genome, estimated at around 3,000 lesions per cell per day in unstressed conditions [9, 10]. If not detected in time by cellular DNA repair mechanisms, that are often inactivated in cancer cells [11], the G→8-oxoG oxidation can lead, after replication, to a G→T base transversion [12], which can then get fixed in the cell’s genome. Other well-known examples of mutational processes are those related to cell exposure to tobacco smoke, which leads to an overrepresentation of C*→*A mutations in lung cancer genomes [13, 14], and to ultraviolet (UV) light, which results in an overrepresentation of C*→*T mutations in melanoma [15].

An attempt at a systematic classification of SBSs and their aetiologies has been made in the COSMIC database [16], where 95 mutational signatures from different cancer types have been identified and linked to distinct mutational processes. Nevertheless, the precise identification of their underlying mechanisms remains highly challenging, and about one-fourth of the COSMIC signatures still have an unknown aetiology [17].

The formation of SBSs has been linked to the presence of electron holes migrating along the nucleobase stacks of the double-stranded DNA molecules that make up the genome [18, 19, 20, 21]. Indeed, the nucleobases, which are the aromatic moieties of the nucleotides arranged in parallel *π*–*π* stacks along each of the two strands, can be easily ionized by various physical and chemical agents.

The resulting electron holes can migrate along the base stack, generally in the 5’ to 3’ direction, without hopping between strands. They remain localized in regions of low ionization potential, sometimes far from their original site of formation [18], and can ultimately trigger SBSs [20, 21, 22, 23, 24].

The DNA ionization propensity and charge migration depend on the nucleobase sequence, with electron holes preferentially accumulating in guanine-rich regions [25, 26]. This preference arises because guanine has the lowest vertical ionization potential (vIP) among the DNA bases [19, 21, 27, 28]. In addition, DNA ionization propensity has been shown to depend, to a lesser extent, on the nucleobase stack conformation [27, 29].

Note that electronic properties of DNA play a fundamental role not only in mutagenesis processes but also in a wide range of physiological processes within the cell. For example, charge transfer appears to be essential in signaling mechanisms to identify and repair DNA damage [30, 31, 32]. Oxidative DNA damage has also been suggested to be involved in gene regulation by functioning as a kind of epigenetic mark [33].

To analyze the relationship between DNA electronic properties and genome mutability, we previously used quantum chemistry approaches to compute the vertical ionization potential (vIP) of stacked nucleobase motifs [27, 29, 34], defined as the energy required to remove an electron from a molecule without taking into account possible structural rearrangements. We showed [34] that DNA motifs with low vIP values are more frequently mutated, whereas those with higher vIP tend to be more protected against mutations.

In this study, we extend the analysis of DNA electronic properties to gain deeper insight into their relationship with cancer mutagenesis processes. Specifically, we compare these properties with experimental data on DNA mutability across both local and genome-wide scales. We further examine the sequence-dependent tendency of DNA motifs to undergo oxidation and its association with the frequency of SBSs across different sequence contexts in somatic mutations from normal and cancer tissues, as well as in germline mutations. Finally, we explore how DNA electronic characteristics correlate with known cancer mutational signatures [16], and identify those in which they play a prominent role.

## 2 Materials and Methods

### 2.1 Datasets

We constructed two datasets of annotated somatic mutations, one in normal tissues and another in cancer tissues, and a third dataset of germline variants, among which benign and disease-causing variants. We only considered SBSs, and overlooked insertions and deletions. We provide below a brief description of the dataset characteristics; additional information is available in Supplementary Sections 1-2:

- *S_SC_* is a dataset of somatic SBSs in cancer tissues that we collected from the COSMIC database [16], limited to nuclear DNA. It contains about 12.5 million SBSs; about 40% are situated in coding regions and the remaining 60% in non-coding regions. Among the coding mutations less than 0.5% are classified as cancer driver mutations.
- *S_SN_* is a dataset of somatic SBSs in normal tissues that we collected and curated from SomaMutDB [35]. It contains about 8 million SBSs, of which only 2.9% are located in coding regions, with the rest occurring in non-coding regions.
- *S_G_* is a collection of germline SBSs from ClinVar [36], which have been carefully annotated with genomic and clinical annotations. After curation, we obtained information for about 3 million germline SBSs, among which 80 % are in coding regions. In total, 36% and 5% of the mutations are classified as benign and pathogenic, respectively, while the remaining 49% are variants of unknown significance or have conflicting interpretations.

We downloaded the variant data in June 2025 and filtered them to retain only SBSs. These variants were then annotated using snpEff (v5.2e) [37] based on the GRCh38 reference genome. They were classified into different categories: coding, separated into nonsense, missense and silent; untranslated region (UTR); splice region found at the beginning and end of the introns and spanning the intron–exon junction; introns; and intergenic (see Supplementary Table 1). Only canonical transcripts were considered.

### 2.2 Sequence context of the variants

For each SBS in each of the three datasets *S_SC_*, *S_SN_* and *S_G_*, we collected the flanking sequence of the mutated base ’X’ in the reference genome, both upstream and downstream. More precisely, we considered motifs with *n* nucleobases upstream and *n* downstream (*n ≤* 5), noted 5’-N..NXN..N-3’, or simply N..NXN..N, with the mutated base X at the middle position and N one of the four nucleobases. The frequency of occurrence of the motifs, *f*_N..NXN..N_, was computed separately in the three mutation datasets. In addition, we computed the frequencies 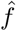_N..NXN..N_ of the same base motifs in the reference human genome GRCh38. The natural logarithm of the normalized frequency of each mutated motif, called *F* , is defined as:

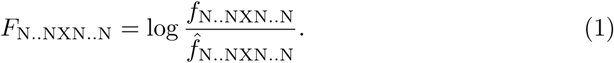

The more negative the *F* value, the less frequently the variant is observed in the N..NXN..N context compared to a random mutation in the genome. In contrast, positive *F* values indicate that the variant is more frequent in that specific nucleotide motif. Note that we took the normalizing frequencies 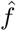 independent of the DNA region, e.g. coding and non-coding, given that the electron hole migration is essentially blind to it. This choice is discussed in the Conclusion section.

Most sequencing datasets do not retain information about the original strand on which a mutation first occurred, since sequencing is performed after the variant has been fixed. Consequently, we calculated the 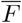 values from the sum of the frequencies of the motifs N..NXN..N and their reverse complement 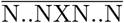 :

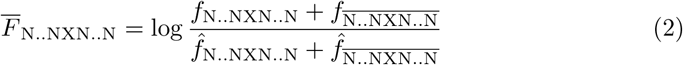

### 2.3 Vertical ionization potentials of nucleobase motifs

The vIP is the energy needed to remove an electron from a molecule, without a change in geometry. Thus, vIP values are always positive, and the lower they are, the easier it is to extract an electron.

In a previous paper [38], we have introduced the freely available program vIPer (github.com/3BioCompBio/vIPer) that computes the vIP value of all possible single-stranded DNA stacks in the commonly observed B conformation. More precisely, we have calculated the vIP values for all single nucleobases N, doublets NN, triplets NNN, and quadruplets NNNN using *ab initio* quantum chemistry methods. For longer nucleobase sequences, we have developed an iterative computational model, called vIPer, which is based on the vIP value of the submotifs included in the motif, and is able to return the vIP values of a motif of any length.

We also calculated the vIP of double-strand motifs, noted 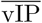, as the mean of the vIP values of both the forward and reverse strand motifs:

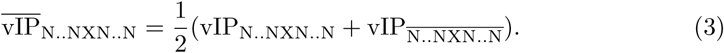

Both vIP and 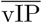 values can directly be obtained from the vIPer program.

Among all four nucleobases, guanine has been experimentally shown to be the nucleobase from which electrons are most easily extracted, and therefore it has the lowest vIP value, followed by adenine, cytosine, and thymine [39]. On the average, longer motifs exhibit lower vIP values; however, the magnitude of this decrease diminishes as motif length increases. For example, the mean vIP computed by vIPer is equal to 8.37, 7.90, 7.68, 7.54, 7.43 eV for singlets, doublets, triplet, quadruplets and quintuplets, respectively [38] (Supplementary Figure 1.a). The vIP values are obviously related to the guanine content of the motifs. This relation is illustrated in Supplementary Figure

1.b for quintuplet motifs. As expected, the correlation is strong but imperfect: two different motifs with the same guanine content can differ in vIP by as much as 1 eV.

### 2.4 DNA oxidative damage and mutational signature

We collected the 95 mutational signatures contained in the COSMIC database [16], each corresponding to a different aetiology. Each signature is characterized by the probability of observing a triplet motif NXN mutated into NYN. By convention, the triplets reported in the signatures are those on the DNA strand in which the mutated nucleobase is a pyrimidine (C or T) [16]. There are thus six possible substitution types: C*→* (A, G, T), and T*→* (A, G, T), and a total of 96 double-stranded triplet motif substitutions NXN*→*NYN for each mutational signature. In our analysis, we only considered the wild-type motifs and not the specific substitutions. We thus took into account the frequency of the 32 wild type triplet motifs and their reverse complements in each signature, divided it by the frequencies 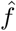 of the same motifs in the reference human genome, and took the logarithm of this ratio, thus obtaining 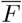_NXN_ defined in Eq. (2).

### 2.5 DNA oxidative damage data

We collected and analyzed published experimental data on DNA oxidative damage at the genome scale, generated by two high-throughput sequencing techniques, each designed to detect distinct chemical signatures of oxidative stress in DNA [40, 41].

The first technique is repair-assisted damage detection (RADD) sequencing, which locates oxidative DNA lesions at about 150 base pair (bp) resolution [41]. DNA was first exposed to the oxidizing agent potassium bromate. Repair enzymes were then used to excise the DNA lesions, and to repair them *in vitro* with biotinylated nucleotides. The positions of the lesions were identified through standard DNA fragmentation, immunoprecipitation, and sequencing [41]. The data were reported on consecutive 200 bp segments of DNA.

The second technique is the apurinic-site (AP) sequencing approach, which is designed to detect apurinic sites induced by X-ray treatment [40]. AP sites are first labeled with an aldehyde-reactive probe, forming a biotin-tagged DNA adduct. This is followed by DNA fragmentation, immunoprecipitation, and sequencing, allowing for genome-wide mapping of DNA lesions at approximately 250 bp resolution. Three replica of this experiment were performed in [40] and the data was provided for genome segments of heterogeneous length.

To enable direct comparison between the AP and RADD data over identical genomic regions, we first computed per-nucleotide AP values for each of the three replicates. For each segment, the strength of the AP signal was divided by the segment length, and the resulting normalized value was assigned to all nucleotides within that segment. The per-nucleotide values were then averaged across replicates. To obtain an AP value for each genome region of 200 bp defined by the RADD data, we simply summed the per-nucleotide values within that region. The 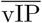 values were computed for all these 200 bp genome segment using the vIPer program.

We considered the AP and RADD data and the vIP values at multiple genomic resolutions: we first considered these data at the finest resolution of 200 bp, and then combined several successive genome regions to form regions of up to 100,000 bp, which is the size proposed in [40] to reach reasonable AP data accuracy. The RADD, AP and vIP values in these larger DNA segments were taken as the average of the scores of the combined segments.

## 3 Results

### 3.1 Genome-scale oxidation damage data and vIP

Over the past few decades, many experimental studies have been devoted to quantifying DNA oxidative damages, from single-site to genome-wide levels, and to improve our understanding of the biophysical processes that cause them. Here, we collected a series of data to investigate more deeply the DNA sequence dependence of oxidative damage and compared them with calculated vIP values, which estimate how easily a DNA motif oxidizes. This is fundamental for understanding the genetic stability of the genome and for potentially identifying mutational hotspots in the genome.

In a previous study [38], we have analyzed single-site–level data on the photoinduced oxidation of twelve TNGNT motifs [42], and demonstrated a strong anticorrelation between measured relative reactivity and calculated vIP values. Here we considered data obtained through two high-throughput techniques developed to study oxidative damage on the genome scale: RADD sequencing of DNA oxidized using potassium bromate [41] and AP sequencing after X-ray treatment [40]. As detailed in

We also calculated vIP scores on these segments. As we do not have information on which strand the mutation primarily occurs, we rather considered the mean of the vIP of both strands, i.e. 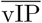 as defined in Eq. (3).

We computed the Spearman correlation between the two experimental oxidation damage scores and the 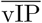 values on the whole human genome, for the original 200 bp segments and aggregated consecutive segments of up to 1,000,000 bp. 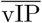 and oxidative damage scores are anticorrelated, as lower vIPs indicate easier electron extraction; we thus correlated the experimental scores with *−*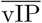.

The results are shown in Figure 1 and Table 1. The correlation improves as a function of segment length, and reaches good values in the 20 kbp to 1,000 kbp range, as already noted for AP replicas [40]. Indeed, there is a certain level of variability in these experiments, as the formation of electron–hole pairs is intrinsically related to probabilistic processes, which hinder accurate mutability estimation at small-scale in such genome-wide experiments.

**Fig. 1:**
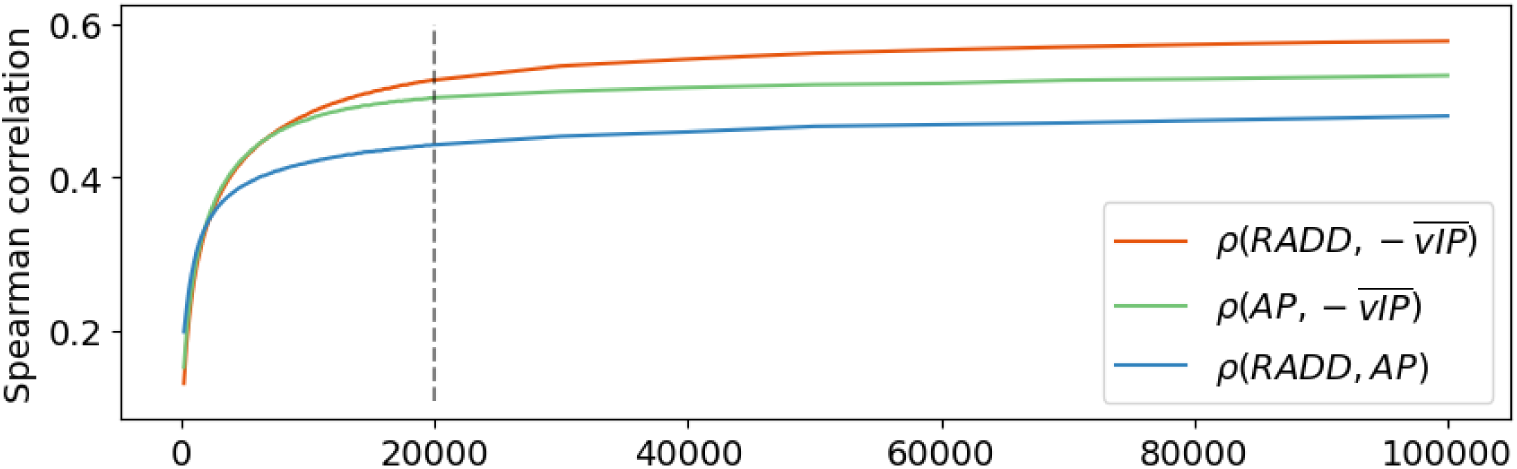
Evolution of the Spearman correlation coefficient *ρ* between RADD, AP and 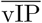 scores as a function of the genome segment length (in number of bp).

**Table 1:** Spearman correlation coefficient *ρ* between RADD, AP and *–*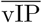 scores for genome segments of various lengths.

| | $-\overline{vIP}/RADD$ | $-\overline{vIP}/AP$ | $RADD/AP$ |
| --- | --- | --- | --- |
| 200 bp | 0.13 | 0.15 | 0.20 |
| 20 kbp | 0.53 | 0.50 | 0.44 |
| 100 kbp | 0.58 | 0.53 | 0.48 |
| 1 Mbp | 0.60 | 0.60 | 0.52 |

The best correlation values are obtained between experimental oxidation scores and *−*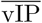 values: 0.53 for RADD scores and 0.50 for AP considering 20 kbp segments, reaching 0.60 for 1 Mbp segments. These values highlight the close agreement between our estimates of the electronic oxidative properties of DNA and the experimental mutagenesis data.

Notably, the correlation between experimental RADD and AP scores is significantly lower, i.e. 0.44 and 0.52 for 20 kbp and 1 Mbp segments, respectively. This likely reflects differences in the mutagenic agents and cell types used in each experimental assay, which result in distinct patterns of DNA accessibility. Specifically, RADD exposes an osteosarcoma cell line to potassium bromate [41] whereas AP exposes the hepatocyte-derived HepG2 cell line to X-ray irradiation [40].

To further investigate the relationship between experimental and computed oxidation DNA lesions, we plotted RADD, AP and (*−*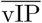) scores along the reference human genome. For comparison purpose, we normalized these scores based on their ranks. The results are shown in Figure 2 for chromosome 1 and in Supplementary Figure 2 for all other chromosomes. We plotted the results using 100 kbp segments to obtain a smooth rendering. We observe that the three curves are consistent and follow each other closely, in all chromosomes, identifying regions with low vIP that are highly mutated.

**Fig. 2:**
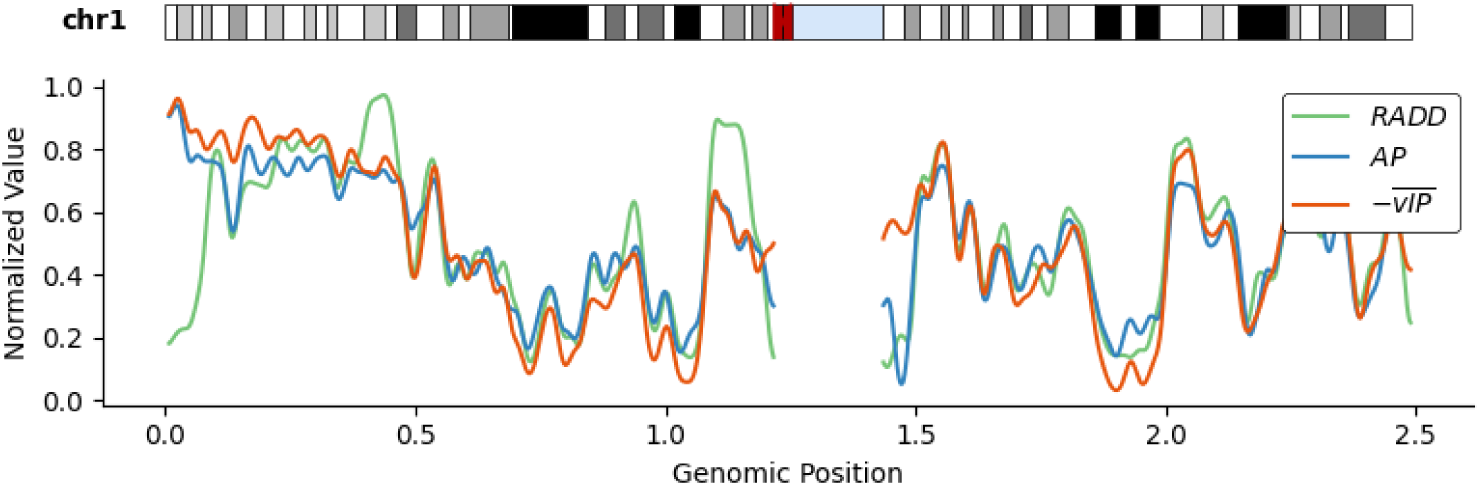
Normalized values of experimental RADD and AP scores and of calculated *−*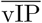 values along human chromosome 1. All the scores were averaged within 100 kbp segments, rank-normalized, and subsequently smoothed using a Gaussian filter spanning ten bins, as implemented in the ndimage.gaussian filter1d function from the *SciPy* Python library. A karyotype of the chromosome is displayed at the top; the blacker regions, the lesser genes and the more compacted; the red region is the centromer. The results on all the other human chromosomes are available in Supplementary Figure 2.

We observe a high variation in mutability across the entire genome, resulting from a complex interplay of genetics and epigenetics [43]. DNA electronic properties play a key role in the mutability process, as shown by the agreement between (*−*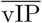) values and experimental mutability scores. However, DNA accessibility is another important factor as already observed in [41]: heterochromatic regions, identified as dark bands in the karyogram, are highly condensed and less accessible to ROS; they generally appear as minima in Figures 2 and Supplementary Figure 2. In contrast, light bands, corresponding to open euchromatic regions, are more prone to mutation.

The (*−*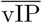) values constitute a very good proxy of mutability, with overall high correlation with the experimental mutability scores. The correlation strength varies within each chromosome, with certain regions displaying almost perfect correlations and others much lower ones. This may stem from local genomic and epigenomic features that are expected to modulate DNA damage, among which DNA methylation, DNA accessibility or alternative DNA structures, which are tissue-dependent and not accounted for in our current model (see Discussion and Conclusion).

### 3.2 Cancer mutational signatures and vIPs

In order to understand the relationship between DNA electronic properties and carcinogenesis, we analyzed the 95 cancer mutational signatures from the COSMIC database [16] and correlated them with the calculated vIP values, as described in the Methods section. Each signature corresponds to a different cancer aetiology. Some are directly related to DNA oxidation processes, such as SBS18, which is associated with ROS, whereas others, such as SBS2 linked to APOBEC-mediated deamination are not.

In a first step, we classified all signatures into six categories, following [44]: mutations resulting from exposure to electrophilic agents, exposure to UV light, defects in DNA repair mechanisms, other mutational processes, experimental artefacts, or unknown causes. Details on this classification is provided in Supplementary Table 4. Note that the identification of mutational signatures in which DNA oxidation plays a major role is a non-trivial issue and clearly depend on the flanking sequence of the mutated bases [33, 45].

To explore this further, we computed, for each signature, the correlation between *F*_NXN_, the logarithm of the normalized mutation frequency of the base triplets NXN defined in Eq. (1), with their calculated vIP values (see Methods). As we do not have information on which strand the mutation primarily occurs, we rather considered the correlation between the averages over both strands, i.e. between 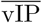 (Eq. (3)) and 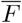 (Eq. (2)). Note that using the vIP of the reference strand or of the strand of minimal vIP still yields strong (although slightly weaker) anticorrelations (see Supplementary Table 5). We considered here the Pearson correlation as the relationship between an energy (vIP) and the logarithm of a frequency (*F* ) is expected to be linear in the context of the Boltzmann law; we added for comparison the Spearman correlation in Supplementary Table 6 and Supplementary Figure 3. Both correlation types show the same trends and lead to the same conclusions.

The results of the 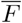 − 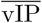 Pearson correlations for the 68 signatures that are not experimental artefacts are shown in Figure 3. We observe that the majority of signatures (72%) show a negative correlation. This proportion is even larger for the signatures categorized as being due to electrophilic substances (80%) or to defects in repair mechanisms (80%). Clearly, only mutations due to oxidative DNA damage can be expected to be related with vIP strength.

**Fig. 3:**
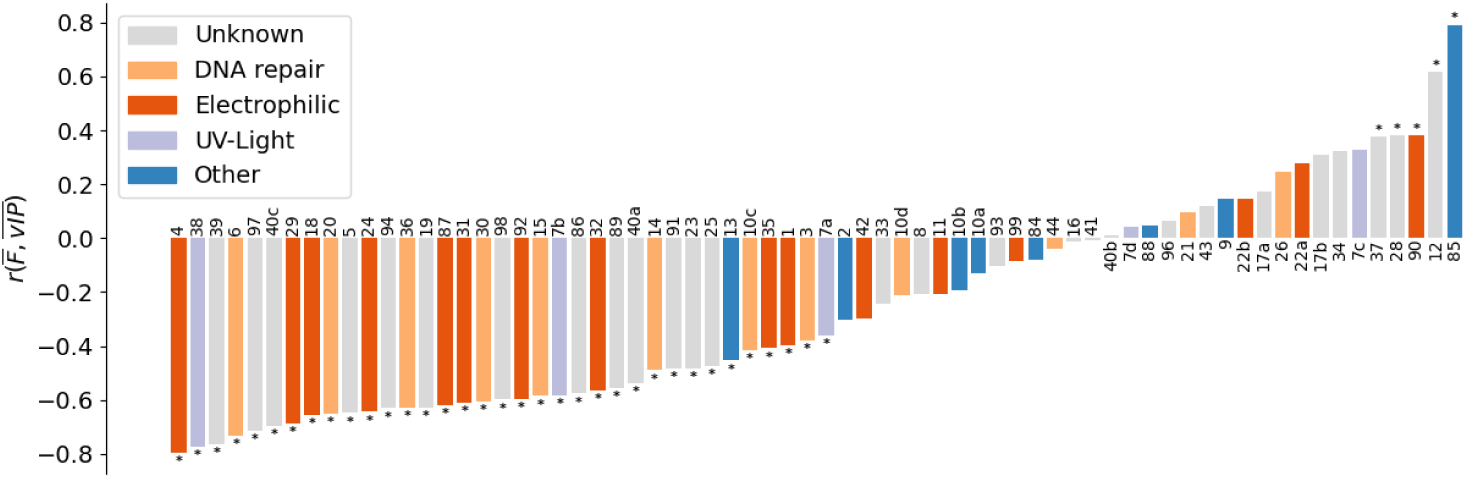
Pearson correlation coefficients between 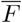 and 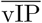 for all cancer mutational signatures that are not reported to be experimental artefacts. The classification is performed according to [44]: exposure to electrophilic agents, exposure to UV light, defects in DNA repair mechanisms, other mutational processes or unknown mechanisms (see Supplementary Table 4). The ’*’ sign indicates that the corresponding p-value is at most 0.05. A similar figure based on Spearman correlations is depicted in Supplementary Figure 3.

The reason why DNA repair–associated mutational signatures are anticorrelated with 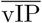 is that several of them, such as the base excision repair–related signatures SBS36 (*r* = *−*0.63) and SBS30 (*r* = *−*0.61), are direcly linked to the repair of oxidative DNA damage. When the repair machinery is impaired, oxidative lesions remain unrepaired and consequently accumulate as mutations of oxidative origin. Mutational signature SBS6 also shows a strong anticorrelation with 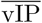 (*r* = *−*0.73). SBS6 is primarily associated with mismatch repair deficiency, often resulting in microsatellite instability, and is responsible for correcting DNA polymerase errors. Despite this, it also correlates strongly with vIP, suggesting that it may additionally be related to oxygen-induced genomic mismatches, as recent evidence appears to indicate [46].

The SBS4 signature shows the strongest anticorrelation: *r* = *−*0.79. It is attributed to tobacco smoking and annotated as likely resulting from DNA lesions directly induced by tobacco carcinogens [44]. Related signatures are SBS92 (*r* = *−*0.60), linked to tobacco smoking, and SBS29 (*r* = *−*0.69), associated with tobacco chewing. The SBS5 signature (*r* = *−*0.65) has an unknown aetiology, yet its mutational burden is annotated to be consistently elevated in cancers related to tobacco exposure. These four signatures show a strong anticorrelation with 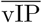, consistent with the presence of a large variety of electrophilic molecules in tobacco, such as the polycyclic aromatic hydrocarbons, that are capable of inducing oxidative DNA damage [47].

Another signature with a strong anticorrelation of –0.65 is SBS18. This is expected, as SBS18 is linked to ROS, which react with DNA to generate a variety of oxidative lesions, (e.g., 8-oxoG). ROS arise both from normal cellular metabolism, especially mitochondrial activity, and from external stressors such as radiation or pollutants, making oxidative DNA damage a burden for the cell [47, 48].

It is also noteworthy that the signature that exhibits the strongest 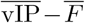 Pearson anticorrelation (*r* = *−*0.81) is SBS45 (Supplementary Table 4); it is not shown in Figure 3 as it is annotated as an artifact. Notably, these mutations arise from the oxidation of guanine to 8-oxoG during sequencing sample preparation, a process driven by oxidative stress [49]. This explains why SBS45 shows such a pronounced 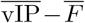 anticorrelation.

Surprisingly, SBS90 shows a positive correlation with 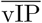 although it is associated with duocarmycin exposure, an electrophilic substance. However, its specific mode of action explains the positive rather than negative correlation: it binds within the minor groove of DNA, where it induces adenine alkylation [50]. Therefore, its specificity for adenine bases severs potential correlation with vIP.

UV radiation induces diverse forms of DNA damage [51]. For example, it can cause oxidative damage either directly, through ionizing interactions with DNA, or indirectly, via UV-induced free radicals and ROS. Conversely, it can also generate double-strand breaks that are not directly linked to oxidative reactivity. Annotations generally do not explicitly specify the underlying mutational processes, but we can reasonably assume that certain UV-related signatures may primarily be due to electron-extracting processes, whereas others may arise from distinct, non-oxidative pathways. The correlation pattern observed with vIP values may therefore help disentangle which UV signatures are predominantly associated with specific types of photoproducts. Although the aetiology of the SBS38 signature is annotated as unknown, it is detected exclusively in UV-associated melanomas and shows a statistically significant correlation of *r* = -0.57, which suggests that it could be caused by indirect DNA oxidation damage caused by UV light. Signatures SBS7b and SBS7a are also due to UV exposure and have statistically significant anticorrelations of *r* = *−*0.58 and *−*0.36, respectively. In contrast, SBS7d and SBS7c display a positive, but not statistically significant correlation, suggesting they might not be due to UV-caused oxidative damage.

### 3.3 Comparison between somatic and germline mutations

Here we analyzed the correlation between the 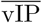 of mutated motifs and their normalized frequency of observation 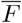 in the datasets of somatic cancer mutations (*S_SC_*), somatic normal tissue mutations (*S_SN_* ), and germline mutations (*S_G_*) (see Methods). The Pearson correlations are shown for both triplets (Table 2.a) and quintuplets (Table 2.b); Spearman correlations are added for comparison in Supplementary Tables 6.a-b. Note that the analysis of germline mutations further elaborates on our earlier studies [34, 38].

**Table 2:**
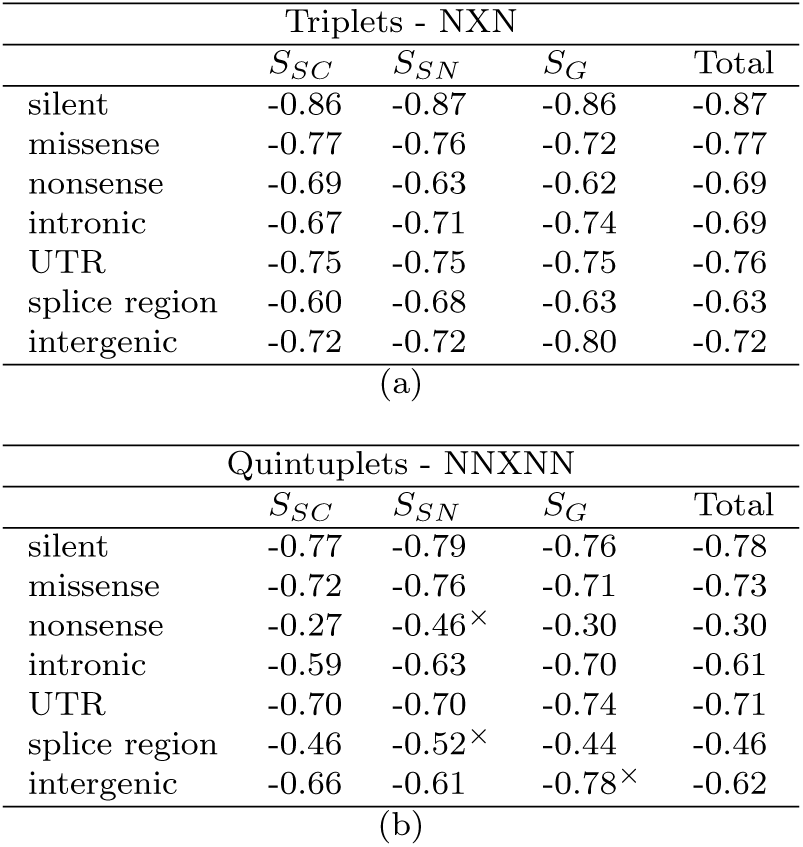
Pearson correlation coefficients *r* between 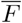 and 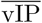 values of (a) triplets and (b) quintuplets in the three datasets *S_SC_*, *S_SN_* , *S_G_*. ”Total” means the ensemble of the three datasets. The p-values are all lower than 0.001. A *×* superscript indicates that the dataset contains less than 50 times the number of triplets (which is 64; in (a)) or of quintuplets (1024; in (b)); the dataset content is specified in Supplementary Table 2. We disregard these values with low statistical significance in our analyses. Similar Tables with Spearman correlations are found in Supplementary Table 6.

First, we observe strong Pearson anticorrelations of –0.73 for triplets and –0.60 for quintuplets when averaged across all datasets and substitution types. These highly significant anticorrelations indicate that lower vIP values are associated with higher mutability. This trend is consistently observed across somatic mutations in both normal and cancer tissues, as well as in germline mutations, with comparable correlation values. Interestingly, this suggests that these mutational processes may arise through similar mechanisms, with oxidative damage playing a central role.

Within coding regions, we classified mutations into silent, missense, and nonsense. Silent mutations show the strongest 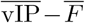 anticorrelations across all three datasets: –0.86 to –0.87 for triplets and –0.76 to –0.79 for quintuplets. Missense mutations also display strong anticorrelations, though to a lesser extent (–0.72 to –0.77 for triplets and –0.71 to –0.76 for quintuplets).

The higher anticorrelations observed for silent relative to missense mutations may arise from their lack of effect on the encoded amino acid, rendering them generally subject to weak selective pressure and resulting in a correlation pattern that more closely reflects the underlying mutational process. Note that silent mutations are not all neutral: they can affect translation and mRNA stability, with some linked to cancer pathogenesis [52, 53, 54]. In contrast, missense mutations are more strongly shaped by purifying selection, which removes many deleterious variants and therefore distorts the underlying 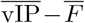 relationship. Nonsense mutations, which introduce premature stop codons, are typically the most deleterious and therefore subject to even stronger selective pressure, resulting in the weakest anticorrelations (–0.62 to –0.69 for triplets and –0.27 to –0.30 for quintuplets). Overall, the decline in anticorrelation from silent to missense to nonsense mutations likely reflects their increasing functional impact and the corresponding intensification of negative selection.

In non-coding regions, the strongest 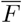 − 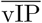 anticorrelation occur in untranslated regions (UTRs), which show anticorrelations comparable to missense mutations (*−*0.75 for triplets and *−*0.70 to *−*0.74 for quintuplets). Indeed, UTRs regulate mRNA stability, localization, and translation, so mutations in these regions can significantly disrupt gene expression [55].

Splice regions show weaker anticorrelations (–0.60 to –0.68 for triplets and –0.45 for quintuplets), consistent with their high functional impact. Mutations in these sites often produce aberrant transcripts and are well known to contribute to inherited diseases and cancer [56, 57].

Introns display intermediate anticorrelations (–0.67 to –0.74 for triplets and –0.59 to –0.70 for quintuplets). Some intronic motifs have been reported to play a role of limiting the impact of oxidative damage. It has indeed be observed that poly-Gua regions are frequent near the 5’ termini of introns, where they have been suggested to serve as sacrificial anodes to protect the protein-coding part of the genes from oxidative damage [58]. Further analyses are needed for interpreting the mechanisms and impact of mutations in these regions.

Finally, the anticorrelations in intergenic regions are in the *−*0.72 to *−*0.80 range for triplets and about *−*0.62 for quintuplets. The observed variability between datasets, with stronger anticorrelations in the *S_G_* set, may stem from differences in DNA sequencing methods. Specifically, capture sequencing and whole-exome sequencing tend to focus on intergenic regions adjacent to genes which are enriched in regulatory motifs, whereas whole-genome sequencing targets the entire genome. *S_G_* essentially contain variants collected with the former methods and *S_SN_* with the latter; *S_SC_* contains both types.

When interpreting differences in anticorrelation values across genomic regions, it is important to recall that *F* is obtained by normalizing the mutated motif frequency by the motif frequency in the whole genome (see Eq. (1)). We made this choice because the electron hole migration towards the DNA region of lowest vIP is blind to the type of DNA region it crosses. However, although the electron holes can migrate over long distances, of at least 100 Å [59], they do not migrate across the whole genome. Moreover, some regions may adopt conformations or interact with proteins which either block or facilitate the migration [60]. So, the chosen normalization is an approximation, and differences in the levels of anticorrelation values in different regions may partly be related to this choice. Note, however, that altering the normalization scheme by computing motif frequencies within specific genome regions, does not substantially affect the results, as discussed in Supplementary Section 8.

*Influence of flanking sequences.* We observed a decrease in 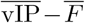 anticorrelation strength between triplets and quintuplets, accompanied by a strong increase in statistical significance, as measured by p-values of about 10*^−^*^10^ and 10*^−^*^100^ for triplets and quintuplets, respectively. Here, we investigated whether this anticorrelation persists for longer motifs, assessing whether flanking sequence context beyond quintuplets and its associated 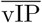 continues to modulate DNA mutability. We specifically focused on silent mutations, where the negative selection is lowest. As shown in Figure 4, the strength of anticorrelations slightly decrease with motif length to approximately the same extent for the three datasets *S_SC_*, *S_SN_* , *S_G_*, but remain strong: about *−*0.5 for undecuplet motifs. In parallel, the statistical significance increases sharply, reaching an undetectable *p*-value for undecuplets. The decrease in the strength of the 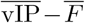 anticorrelation may be attributed to the increasing uncertainty in vIP calculations for longer motifs and the limited availability of mutational data to adequately sample all motifs.

**Fig. 4:**
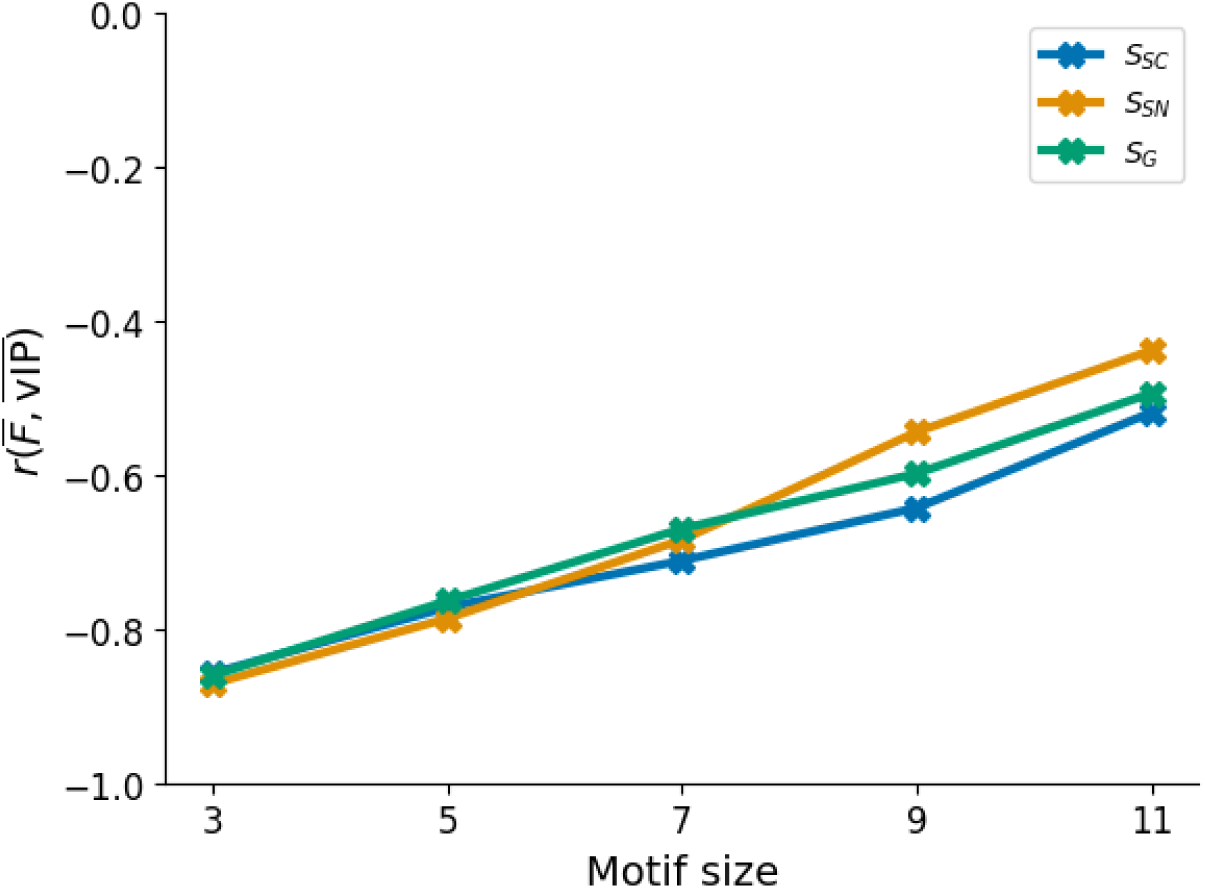
Pearson correlation between 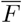 and 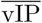 as a function of motif length, for silent mutations in the datasets *S_SC_*, *S_SN_* , *S_G_*.

However, the persistence of significant anticorrelation values underscores the important influence of flanking sequences on both vIP and mutation frequency. This result strongly supports the causative link between vIP and oxidative, mutation-inducing phenomena, which are both non-local in nature. Indeed, oxidative damage has been experimentally shown to occur sometimes far from the eventual mutation site, and electron holes can migrate along the DNA stack over long distances to reach regions of lowest vIP [59].

*Dependence on chromosome.* We tested whether the strength of the anticorrelation between 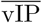 and 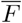 depends on the chromosome. For this purpose, we considered together all the silent mutations contained in the datasets *S_SC_*, *S_SN_* , *S_G_*. As shown in Supplementary Table 7, there are only minor variations in Pearson anticorrelation across the different chromosomes: from *−*0.73 for chromosome 13 to *−*0.79 for chromosomes 16, 20, and 22. The only exception is the Y chromosome, which shows *r* = *−*0.64. As the most compact and densely packed chromosome, it may be partially ”protected” from oxidative damage. However, the limited number of silent mutations on the Y chromosome (446) precludes this result from reaching statistical significance.

*Dependence on the tissue.* As various tissues encounter diverse levels and types of mutagens and mutation processes, face unique functional constraints, and develop cancers through different molecular pathways, we investigated the 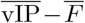 anticorrelation strength of cancer mutations in *S_SC_* according to the tissue; we focused again on silent mutations.

We found the anticorrelations for triplet motifs to lie between *r* = *−*0.87 for mutations in the lung to *r* = *−*0.72 for mutations in the nervous system (Supplementary Table 8). These correlation differences are related to the mutational signatures discussed in the previous section. For example, lung cancer is predominantly caused by oxidative substances present in tobacco, and thus both tobacco-related signatures and mutations in lung anticorrelate strongly with 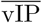. This reflects the dominant contribution of oxidative agents present in tobacco to lung carinogenesis. In contrast, only a subset of DNA lesions caused by UV light are of oxidative nature. As a consequence, different UV-related signatures have different anticorrelation strengths, which can be related to the observation that skin cancer mutations anticorrelate less strongly (*r* = 0.76).

## 4 Discussion and Conclusion

Oxidative DNA damage, generated by endogenous metabolic by-products or exogenous environmental stressors, produces lesions that are key drivers of carcinogenesis and determinants of chemotherapy efficacy. However, gaining a precise understanding of how oxidative agents give rise to SBSs remains challenging, due to the complexity and diversity of oxidative lesions and their repair pathways.

In this study, we advanced this question by studying the DNA electronic properties quantified through 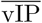 values. Low-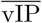 genomic regions are particularly susceptible to oxidative lesions, as they have a greater propensity for electron extraction leading to electron-hole formation. Such lesions are either repaired by the cellular repair machinery or cause sequence alterations such as SBSs. Moreover, when electron holes are generated in higher 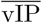 regions, they tend to migrate along the nucleobase stack in the 5*^′^*–3*^′^*direction until they reach a low-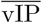 site [61, 62], where a lesion and possibly a mutation may be induced.

In our study, we compared 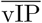 profiles with cancer mutational signatures and demonstrated that such profiles reliably highlight signatures linked to oxidative damage aetiologies. Therefore, our 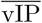-based approach could be used to predict the oxidative nature of cancer mutational signatures of unknown origin, thereby providing new insights into their aetiology. A key perspective of this study involves designing a tool to enable such 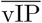-based predictions.

Furthermore, we found that negative selection shapes the mutation spectrum observed in sequencing data. Highly deleterious SBSs, such as those disrupting essential regulatory elements or introducing nonsense mutations, are removed from the cell population and therefore go undetected. This depletion of highly deleterious variants likely explains why severe mutation types (e.g., nonsense and, to a lesser extent, missense) tend to show weaker anticorrelations with vIP than more benign ones such as silent mutations.

An interesting finding is that the 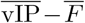 anticorrelations we observed are very similar for somatic and germline mutations, as well as in cancer and normal tissues. This suggests that these mutations arise through similar mechanisms, in which oxidative damage plays a central role. Some small differences are nevertheless observed, which could partly reflect varying levels of negative selection [63, 64].

Although our vIP model already describes DNA electronic properties and their relationship to mutational processes very well, its ability to identify genomic regions prone to mutation could still be improved. Obtaining more precise vIP values using advanced quantum chemistry techniques and considering alternative DNA conformations, thereby dropping the assumption that chromosomes adopt a uniform B-form DNA structure, could further refine the model and provide a more accurate representation of sequence-dependent mutational susceptibility.

While oxidative damage is strand specific, this information is often unavailable in mutation databases, leading us to use the vIP and mutation frequency averaged over both strands. Using the vIP of the exact strand where the oxidative damage occurred could further improve our understanding of these processes. Interestingly, recent results suggest a strand bias in oxidative damage, with transcribed strands experiencing less oxidative damage than their non-transcribed counterparts [45].

It would also be interesting to incorporate chromatin state and tissue-specific genomic data in our model. Chromatin accessibility, nucleosome positioning, and tissue-dependent epigenetic landscapes all influence how DNA is exposed to damaging agents and how efficiently lesions are repaired. Possible important protein-DNA interactions likely to affect electron hole migration [60, 65] could also be integrated. Including these layers of information could therefore improve the ability of our vIP model to describe mutational processes.

## Supporting information

Suppementary Info

## Acknowledgments

This work was supported by a Téévie grant and a PDR grant from the Belgian National Fund for Scientific Research (F.R.S.-FNRS).

