## Supplementary material for "Linking somatic mutations in cancer to the electronic properties of DNA": Suppementary Info

#### Contents

**1 Variant data classification** **2**

**2 Dataset content** **2**

**3 Calculated vIP values of nucleobase motifs** **4**

**4 Chromosome-scale oxidation damage data and vIP** **5**

**5 Mutational Signatures** **7**

**6 Strand dependence of vIP values** **9**

**7 Spearman correlations for somatic and germline mutations** **10**

**8 Comparison with previous analyses** **11**

**9 $\overline{\text{vIP}} - \overline{F}$ correlations per chromosome** **12**

**10 $\overline{\text{vIP}} - \overline{F}$ correlations according to the tissue** **13**

### 1 Variant data classification

We downloaded the data in June 2025 from COSMIC, ClinVar and SomaMutDB and filtered the variants to retain only the single base substitutions (SBS). These variants were annotated using snpEff (v5.2e) [1] based on the GRCh38.mane.1.2.refseq human reference genome. The variants were classified into seven mutually exclusive categories: silent, missense, nonsense, splice region, intronic, untranslated regions (UTR) and intergenic. The classification was performed based on the annotation mapping of Table S1 and the annotations of the ANN INFO field added by snpEff in the VCF files.

| | |
|---|---|
| start_lost | missense |
| upstream_gene_variant | intergenic |
| synonymous_variant | silent |
| 5_prime_UTR_premature_start_codon_gain_variant | UTR |
| stop_lost | missense |
| downstream_gene_variant | intergenic |
| start_retained_variant | silent |
| stop_retained_variant | silent |
| 3_prime_UTR_variant | UTR |
| intron_variant | intronic |
| stop_gained | nonsense |
| missense_variant | missense |
| splice_region_variant | splice_region |
| 5_prime_UTR_variant | UTR |
| intragenic_variant | NaN |
| initiator_codon_variant | missense |
| intergenic_region | intergenic |

Table S1: Mapping of snpEff mutation types [1] to the broader mutation categories used in this study.

### 2 Dataset content

| **type** | $S_{SC}$ | | $S_{SN}$ | | $S_G$ | |
|---|---|---|---|---|---|---|
| **silent** | 1,182,111 | ( 9.5%) | 67,160 | ( 0.8%) | 672,344 | (20.8%) |
| **missense** | 3,317,016 | (26.5%) | 155,487 | ( 1.9%) | 1,875,621 | (58.1%) |
| **nonsense** | 241,261 | ( 1.9%) | 10,669 | ( 0.1%) | 75,732 | ( 2.4%) |
| **intronic** | 6,637,368 | (53.0%) | 2,471,401 | (30.5%) | 396,305 | (12.3%) |
| **UTR** | 495,573 | ( 4.0%) | 86,523 | ( 1.1%) | 69,485 | ( 2.2%) |
| **splice_region** | 70,400 | ( 0.5%) | 7,588 | ( 0.1%) | 116,357 | ( 3.6%) |
| **intergenic** | 564,944 | ( 4.5%) | 5,306,820 | (65.4%) | 23,138 | ( 0.7%) |
| **TOTAL** | 12,510,810 | | 8,110,990 | | 3,228,982 | |

Table S2: Number and percentage of SBSs per dataset and mutation type

| Tissues | Intronic | Intergenic | Splice Region | Silent | UTR | Missense | Nonsense | TOTAL | MIN |
|---|---|---|---|---|---|---|---|---|---|
| skin | 145310 | 25525 | 12906 | 321554 | 40585 | 717527 | 55433 | 1318840 | 12906 |
| large_intestine | 240398 | 20277 | 9223 | 211099 | 59025 | 564009 | 36954 | 1140985 | 9223 |
| liver | 1403416 | 110040 | 8361 | 64131 | 61422 | 171939 | 11101 | 1830410 | 8361 |
| lung | 152018 | 16395 | 7591 | 159388 | 39144 | 520866 | 39774 | 935176 | 7591 |
| stomach | 225310 | 17863 | 6478 | 105626 | 24556 | 296469 | 15566 | 691868 | 6478 |
| haematopoietic_and_lymphoid_tissue | 763685 | 59859 | 5467 | 48538 | 35413 | 172649 | 9952 | 1095563 | 5467 |
| breast | 808755 | 68539 | 4103 | 49676 | 36987 | 134215 | 10444 | 1112719 | 4103 |
| central_nervous_system | 70160 | 7736 | 2537 | 34638 | 10698 | 153233 | 6653 | 285655 | 2537 |
| biliary_tract | 68753 | 5044 | 2394 | 17591 | 5707 | 64659 | 4207 | 168355 | 2394 |
| urinary_tract | 60973 | 7019 | 1904 | 60248 | 12658 | 130118 | 9701 | 282621 | 1904 |
| pancreas | 796480 | 65779 | 1893 | 18087 | 23214 | 61433 | 3998 | 970884 | 1893 |
| small_intestine | 48933 | 2250 | 2060 | 14532 | 4405 | 32526 | 1759 | 106465 | 1759 |
| prostate | 515242 | 43598 | 1733 | 25590 | 20072 | 69528 | 4242 | 680005 | 1733 |
| endometrium | 68586 | 10614 | 1667 | 102014 | 70440 | 283623 | 25521 | 562465 | 1667 |
| oesophagus | 612069 | 51833 | 1561 | 32347 | 18556 | 86488 | 6220 | 809074 | 1561 |
| upper_aerodigestive_tract | 69319 | 7312 | 1317 | 47725 | 9076 | 129987 | 9135 | 273871 | 1317 |
| ovary | 285889 | 24852 | 903 | 15927 | 12138 | 46501 | 2926 | 389136 | 903 |
| kidney | 184693 | 15431 | 764 | 19035 | 10743 | 70699 | 4842 | 306207 | 764 |
| cervix | 4714 | 1507 | 732 | 17167 | 13546 | 43037 | 3951 | 84654 | 732 |
| soft_tissue | 157679 | 14128 | 488 | 8343 | 10026 | 48402 | 1746 | 240812 | 488 |
| thyroid | 93268 | 9267 | 414 | 14541 | 4943 | 80753 | 5201 | 208387 | 414 |
| autonomic_ganglia | 1937 | 408 | 445 | 2761 | 1379 | 14623 | 1028 | 22581 | 408 |
| bone | 12658 | 1244 | 263 | 7898 | 1872 | 14022 | 682 | 38639 | 263 |
| nervous_system | 8487 | 1503 | 219 | 12633 | 984 | 53881 | 3148 | 80855 | 219 |
| meninges | 11230 | 811 | 243 | 1208 | 617 | 3042 | 176 | 17327 | 176 |
| eye | 2052 | 189 | 85 | 420 | 172 | 1984 | 130 | 5032 | 85 |
| adrenal_gland | 220 | 70 | 136 | 642 | 233 | 4911 | 433 | 6645 | 70 |
| parathyroid | 208 | 66 | 69 | 151 | 47 | 5705 | 326 | 6572 | 47 |
| salivary_gland | 685 | 129 | 23 | 1034 | 60 | 7542 | 502 | 9975 | 23 |
| testis | 1187 | 100 | 16 | 435 | 123 | 1209 | 74 | 3144 | 16 |
| peritoneum | 68 | 21 | 14 | 366 | 10 | 1651 | 100 | 2230 | 10 |
| pituitary | 162 | 9 | 17 | 194 | 17 | 639 | 40 | 1078 | 9 |
| placenta | 315 | 107 | 8 | 28 | 29 | 9849 | 298 | 10634 | 8 |
| thymus | 39 | 10 | 311 | 5 | 1342 | 83 | 1790 | 5 | |
| pleura | 89 | 46 | 3 | 253 | 9 | 3586 | 225 | 4211 | 3 |
| genital_tract | 87 | 13 | 1 | 300 | 5 | 1504 | 56 | 1966 | 1 |
| vulva | 31 | 18 | 1 | 882 | 6 | 1879 | 138 | 2955 | 1 |
| lymph_node | 799 | 60 | 8 | 31 | 18 | 1 | 917 | 1 | |
| penis | 11 | 2 | 1 | 1 | 347 | 20 | 382 | 1 | |
| gastrointestinal_tract | 2 | 2 | 1 | 17 | 1 | 23 | 1 | | |
| fallopian_tube | 1 | 5 | 101 | 5 | 112 | 1 | | | |
| vagina | 2 | 23 | 1 | 26 | 1 | | | | |

Table S3: Number of mutations observed in each primary tissue type in the $S_{SC}$ dataset of cancer mutations, stratified by mutation category, reported in in COSMIC [2]. Only mutations detected in tumors originating from the indicated tissue are included. Each SBS in $S_{SC}$ may be observed across multiple tissue types. For each SBS, we assigned as primary tissues all tissue types in which it was observed in primary tumors, excluding metastatic occurrences. The "MIN" column indicates the number of mutations in the least frequent mutation category within each tissue.

#### 3 Calculated vIP values of nucleobase motifs

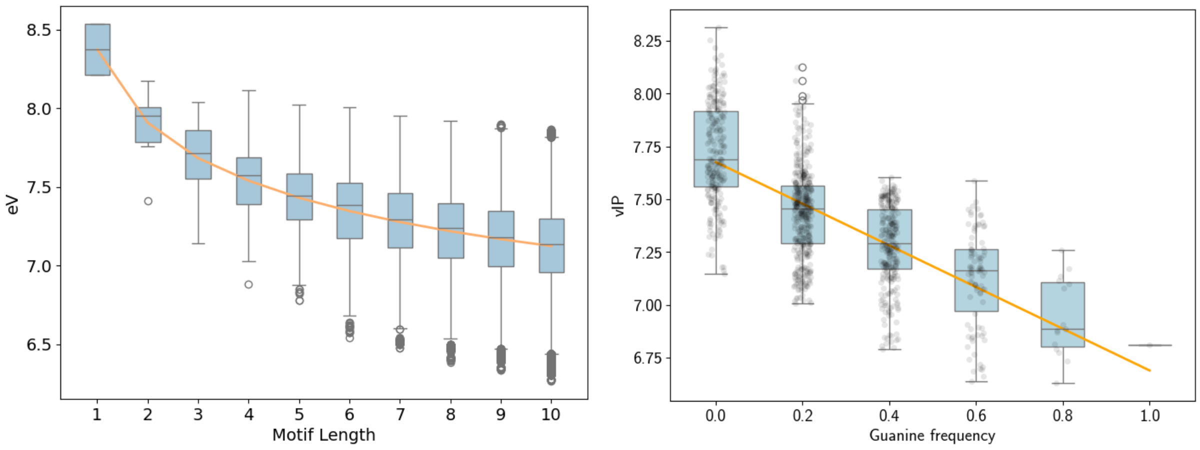


(a) (b)

Figure S1: vIP values of nucleobase motifs, calculated using vIPer [3]. (a) vIP values (in eV) of all possible nucleobase motifs as a function of motif length; (b) vIP values of all possible nucleobase quintuplets as a function of the guanine frequency in the quintuplets.

### 4 Chromosome-scale oxidation damage data and vIP

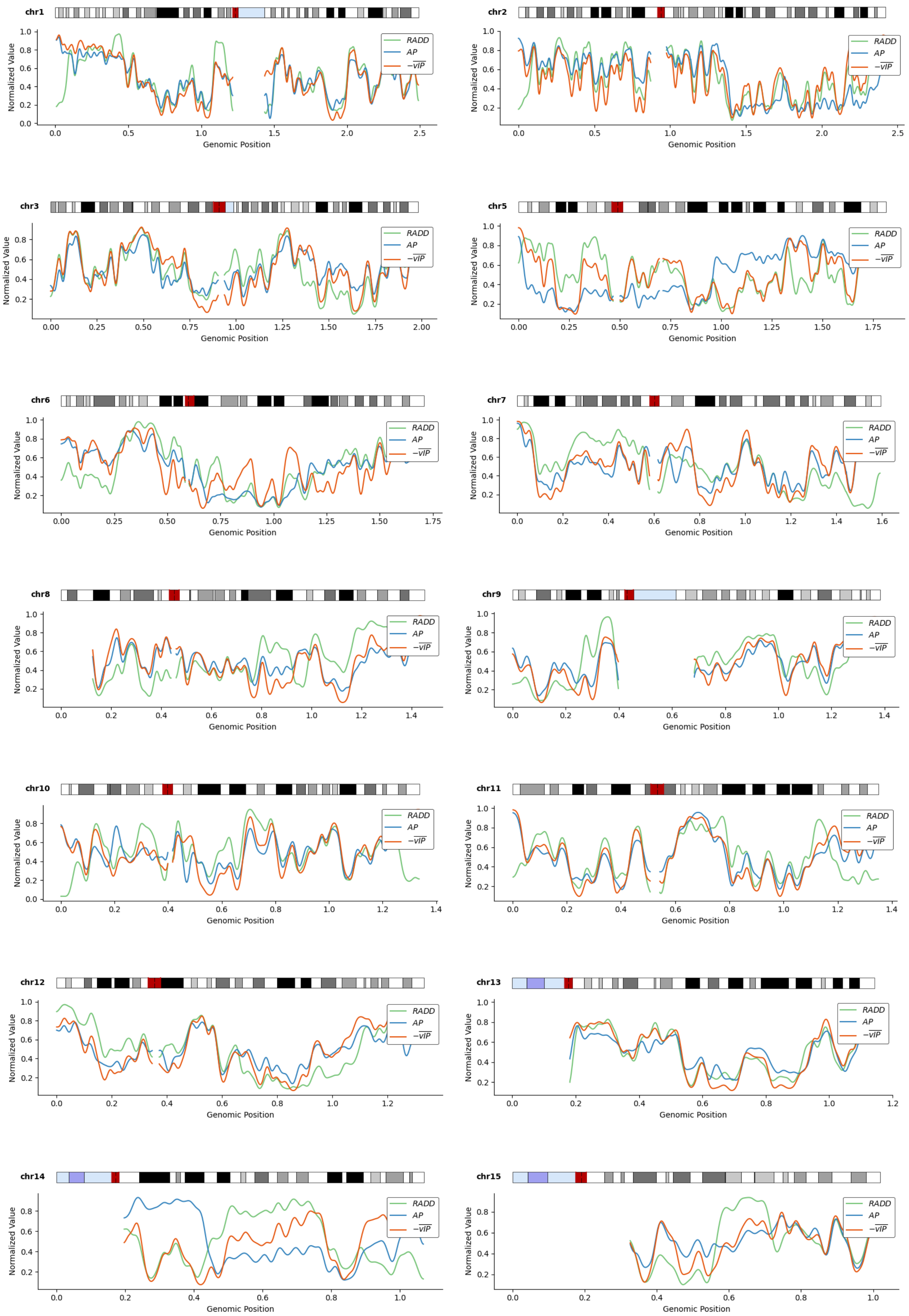

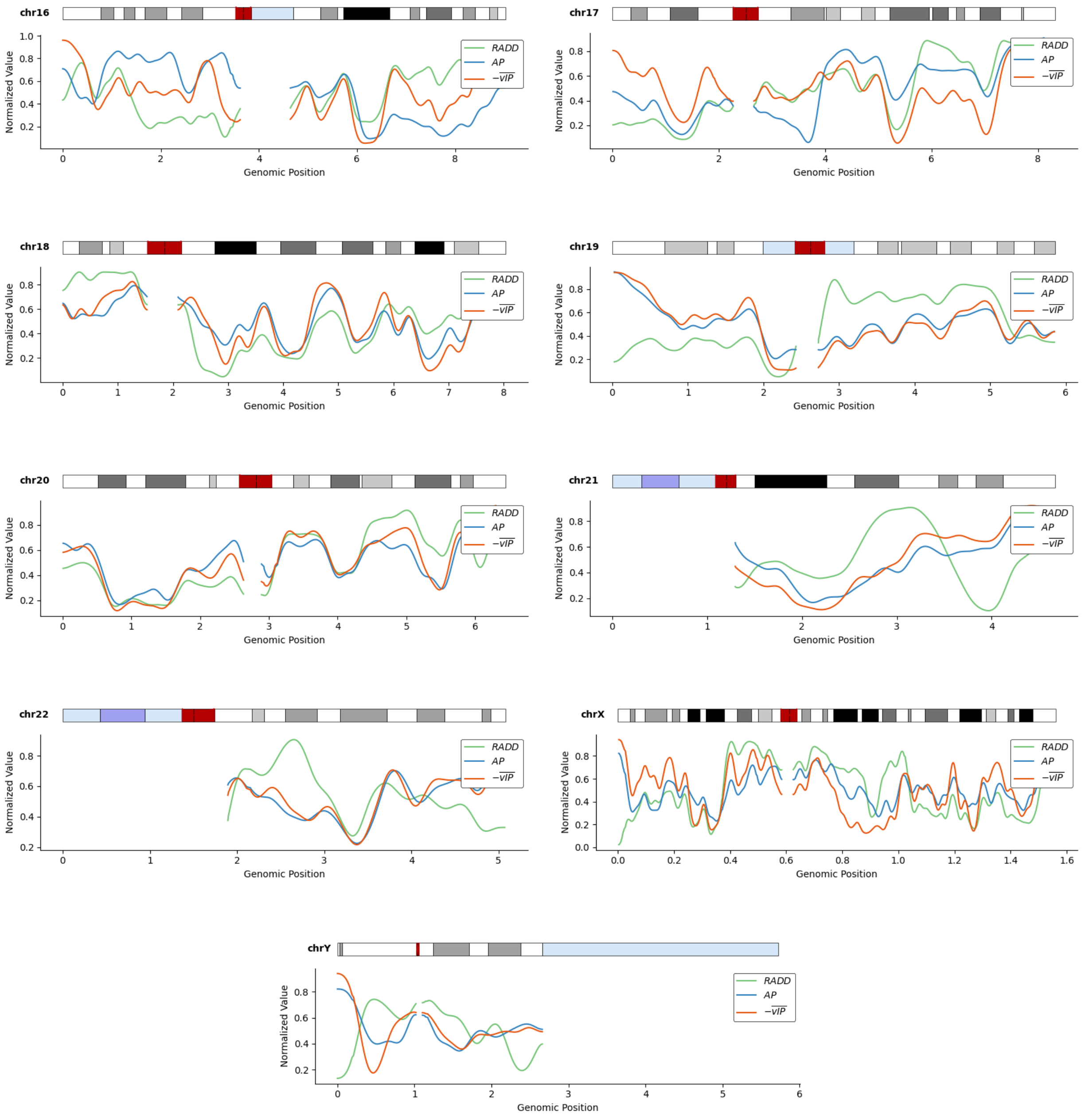


Figure S2: Normalized values of experimental RADD and AP scores and of calculated $-\overline{\text{vIP}}$ values along all human chromosomes. All the scores were averaged within 100 kbp segments, rank-normalized, and subsequently smoothed using a Gaussian filter spanning ten bins, as implemented in the ndimage.gaussian_filter1d function from the *SciPy* Python library. A karyotype of the chromosome is displayed at the top; the blacker regions, the lesser genes and the more compacted; red regions are centromers; purple regions contain tandem repeats of ribosomal RNA genes; light-blue regions are highly polymorphic heterochromatic regions. The results on chromosome 1 is also shown in Figure 2 of the main manuscript.

### 5 Mutational Signatures

| Signature | Proposed aetiology | Classification | $r$ | $\rho$ |
|---|---|---|---|---|
| SBS1 | Endogenous mutational process by deamination of 5-methylcytosine | Electrophilic | -0.40 (2.4e-02) | -0.02 (9.2e-01) |
| SBS2 | Activity of AID/APOBEC family cytidine deaminases. | Other | -0.30 (9.4e-02) | -0.20 (2.7e-01) |
| SBS3 | Defective homologous recombination-based DNA damage repair. | DNA repair | -0.38 (3.3e-02) | -0.38 (3.0e-02) |
| SBS4 | Associated with tobacco smoking. Its profile is similar to the mutational spectrum observed in experimental systems exposed to tobacco carcinogens such as benzo[a]pyrene. | Electrophilic | -0.79 (5.9e-08) | -0.77 (2.7e-07) |
| SBS5 | Unknown. Increased with ERCC2 mutations and tobacco smoking. | Unknown | -0.65 (6.0e-05) | -0.52 (2.2e-03) |
| SBS6 | Defective DNA mismatch repair in microsatellite unstable tumours. | DNA repair | -0.73 (1.9e-06) | -0.72 (3.3e-06) |
| SBS7a | Exposure to ultraviolet light (UV photoproducts). | UV-Light | -0.36 (4.2e-02) | -0.22 (2.2e-01) |
| SBS7b | Exposure to ultraviolet light (UV photoproducts). | UV-Light | -0.58 (4.6e-04) | -0.53 (1.9e-03) |
| SBS7c | UV light. Translesion DNA synthesis opposite photodimers. | UV-Light | 0.33 (6.6e-02) | 0.36 (4.6e-02) |
| SBS7d | UV light. Translesion DNA synthesis opposite photodimers. | UV-Light | 0.04 (8.2e-01) | 0.02 (9.1e-01) |
| SBS8 | Unknown. | Unknown | -0.21 (2.5e-01) | -0.30 (1.0e-01) |
| SBS9 | Polymerase eta during somatic hypermutation in lymphoid cells. | Other | 0.14 (4.3e-01) | 0.18 (3.2e-01) |
| SBS10a | Polymerase epsilon exonuclease domain mutations. | Other | -0.13 (4.7e-01) | -0.16 (3.9e-01) |
| SBS10b | Polymerase epsilon exonuclease domain mutations. | Other | -0.19 (2.9e-01) | -0.08 (6.6e-01) |
| SBS10c | Defective POLD1 proofreading. | DNA repair | -0.42 (1.8e-02) | -0.27 (1.3e-01) |
| SBS10d | Adenoma from individuals with germlinePOLD1exonuclease domain mutations. | DNA repair | -0.21 (2.5e-01) | -0.18 (3.2e-01) |
| SBS11 | Alkylating agent temozolomide treatment. | Electrophilic | -0.21 (2.5e-01) | -0.27 (1.4e-01) |
| SBS12 | Unknown. | Unknown | 0.62 (1.7e-04) | 0.58 (4.4e-04) |
| SBS13 | Activity of AID/APOBEC family cytidine deaminases (APOBEC3A/3B). | Other | -0.45 (9.1e-03) | -0.52 (2.1e-03) |
| SBS14 | Concurrent polymerase epsilon mutation and defective DNA mismatch repair. | DNA repair | -0.49 (4.5e-03) | -0.46 (8.8e-03) |
| SBS15 | Defective DNA mismatch repair. | DNA repair | -0.59 (4.3e-04) | -0.52 (2.5e-03) |
| SBS16 | Unknown. | Unknown | -0.01 (9.5e-01) | -0.18 (3.2e-01) |
| SBS17a | Unknown. | Unknown | 0.18 (3.4e-01) | 0.20 (2.8e-01) |
| SBS17b | Unknown in general. Sometimes, possibly due to fluorouracil chemotherapy or reactive oxygen species. | Unknown | 0.31 (8.3e-02) | 0.29 (1.0e-01) |
| SBS18 | Possibly damage by reactive oxygen species. | Electrophilic | -0.65 (5.0e-05) | -0.65 (5.8e-05) |
| SBS19 | Unknown. | Unknown | -0.63 (1.2e-04) | -0.56 (8.6e-04) |
| SBS20 | Concurrent POLD1 mutations and defective DNA mismatch repair. | DNA repair | -0.65 (5.4e-05) | -0.62 (1.5e-04) |
| SBS21 | DNA mismatch repair deficiency. | DNA repair | 0.10 (6.0e-01) | 0.13 (4.8e-01) |
| SBS22a | Aristolochic acid exposure. | Electrophilic | 0.28 (1.2e-01) | 0.30 (9.0e-02) |
| SBS22b | Aristolochic acid exposure. | Electrophilic | 0.14 (4.3e-01) | 0.09 (6.1e-01) |
| SBS23 | Unknown. | Unknown | -0.48 (5.2e-03) | -0.70 (7.7e-06) |
| SBS24 | Aflatoxin exposure. | Electrophilic | -0.64 (6.9e-05) | -0.65 (4.9e-05) |
| SBS25 | Possibly chemotherapy treatment. | Unknown | -0.47 (6.2e-03) | -0.38 (3.0e-02) |
| SBS26 | Defective DNA mismatch repair. | DNA repair | 0.25 (1.8e-01) | 0.31 (8.2e-02) |
| SBS28 | Unknown. | Unknown | 0.38 (3.1e-02) | 0.45 (9.7e-03) |
| SBS29 | Found in cancer samples from individuals with a tobacco chewing habit. | Electrophilic | -0.69 (1.3e-05) | -0.69 (1.5e-05) |
| SBS30 | Deficiency in base excision repair due to inactivating mutations in NTHL1. | DNA repair | -0.61 (2.3e-04) | -0.55 (1.0e-03) |
| SBS31 | Prior chemotherapy treatment with platinum drugs. | Electrophilic | -0.61 (2.1e-04) | -0.50 (3.3e-03) |
| SBS32 | Azathioprine treatment. | Electrophilic | -0.57 (7.4e-04) | -0.61 (2.1e-04) |
| SBS33 | Unknown. | Unknown | -0.24 (1.8e-01) | -0.28 (1.2e-01) |
| SBS34 | Unknown. | Unknown | 0.32 (7.1e-02) | 0.36 (4.3e-02) |

*Continued on next page*

Table S4 – *Continued from previous page*

| **Signature** | **Proposed aetiology** | **Classification** | $r$ | $\rho$ |
|---|---|---|---|---|
| SBS35 | Chemotherapy treatment with platinum drugs. | Electrophilic | -0.41 (2.0e-02) | -0.37 (3.6e-02) |
| SBS36 | Defective base excision repair, including DNA damage due to reactive oxygen species. | DNA repair | -0.63 (1.2e-04) | -0.56 (8.2e-04) |
| SBS37 | Unknown. | Unknown | 0.38 (3.3e-02) | 0.43 (1.5e-02) |
| SBS38 | Unknown. Found only in ultraviolet light associated melanomas suggesting potential indirect damage from UV-light. | UV-Light | -0.77 (1.9e-07) | -0.70 (9.1e-06) |
| SBS39 | Unknown. | Unknown | -0.77 (3.3e-07) | -0.70 (9.8e-06) |
| SBS40a | Unknown. | Unknown | -0.54 (1.5e-03) | -0.47 (7.2e-03) |
| SBS40b | Unknown. | Unknown | 0.01 (9.6e-01) | 0.10 (5.7e-01) |
| SBS40c | Unknown. | Unknown | -0.70 (9.8e-06) | -0.60 (2.5e-04) |
| SBS41 | Unknown. | Unknown | -0.01 (9.6e-01) | 0.04 (8.1e-01) |
| SBS42 | Occupational exposure to haloalkanes. | Electrophilic | -0.30 (9.7e-02) | -0.35 (4.8e-02) |
| SBS43 | Unknown. Possible sequencing artefact. | Unknown | 0.12 (5.1e-01) | 0.10 (6.0e-01) |
| SBS44 | Defective DNA mismatch repair. | DNA repair | -0.04 (8.4e-01) | -0.33 (6.5e-02) |
| SBS84 | Activity of activation-induced cytidine deaminase (AID). | Other | -0.08 (6.6e-01) | -0.16 (3.8e-01) |
| SBS85 | Indirect effects of activation-induced cytidine deaminase (AID). | Other | 0.79 (6.6e-08) | 0.75 (6.9e-07) |
| SBS86 | Unknown chemotherapy treatment. | Unknown | -0.58 (5.6e-04) | -0.52 (2.3e-03) |
| SBS87 | Thiopurine chemotherapy treatment. | Electrophilic | -0.62 (1.7e-04) | -0.34 (6.0e-02) |
| SBS88 | Exposure to E.coli bacteria carrying pks pathogenicity island, producing genotoxic compound colibactin | Other | 0.05 (8.1e-01) | 0.26 (1.5e-01) |
| SBS89 | Unknown. | Unknown | -0.56 (9.6e-04) | -0.52 (2.4e-03) |
| SBS90 | Duocarmycin exposure. | Electrophilic | 0.38 (3.0e-02) | 0.24 (1.8e-01) |
| SBS91 | Unknown. | Unknown | -0.48 (5.0e-03) | -0.54 (1.5e-03) |
| SBS92 | Associated with tobacco smoking. | Electrophilic | -0.60 (3.2e-04) | -0.54 (1.3e-03) |
| SBS93 | Unknown. | Unknown | -0.10 (5.8e-01) | 0.03 (8.6e-01) |
| SBS94 | Unknown. | Unknown | -0.63 (1.1e-04) | -0.54 (1.5e-03) |
| SBS96 | Unknown. | Unknown | 0.07 (7.2e-01) | 0.16 (3.8e-01) |
| SBS97 | Unknown. | Unknown | -0.71 (4.6e-06) | -0.60 (2.5e-04) |
| SBS98 | Unknown. | Unknown | -0.60 (3.0e-04) | -0.27 (1.4e-01) |
| SBS99 | Melphalan treatment. | Electrophilic | -0.09 (6.4e-01) | -0.10 (5.9e-01) |

Table S4: Mutational signatures from COSMIC [2]: annotated aetiology, classification and Pearson correlation coefficients $r$ with $\overline{\text{vIP}}$ values as well as the corresponding Spearman correlation coefficients $\rho$; p-values are in parentheses.

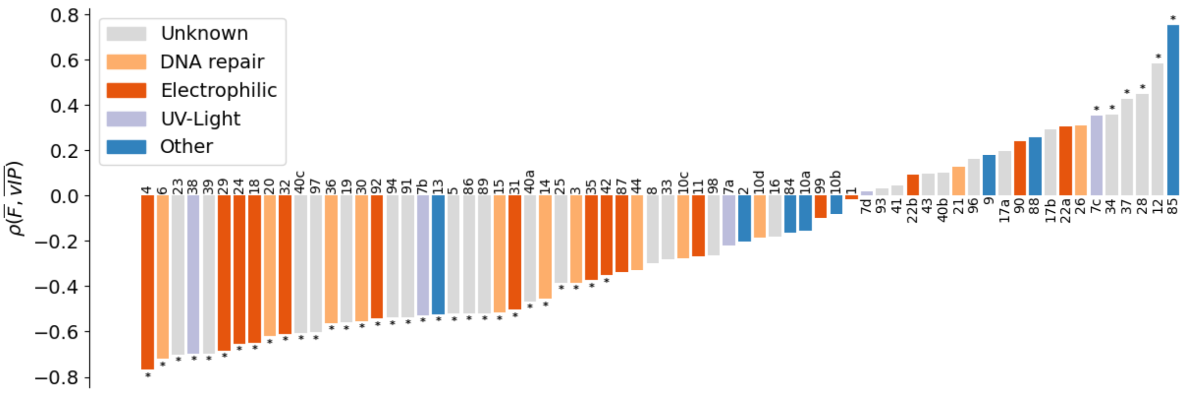


Figure S3: Spearman correlation coefficients between $\overline{F}$ and $\overline{\text{vIP}}$ for all cancer mutational signatures that are not reported to be experimental artefacts. The classification is performed according to [4]: exposure to electrophilic agents, exposure to UV light, defects in DNA repair mechanisms, other mutational processes or unknown mechanisms (see Supplementary Table S4). The '*' sign indicates that the corresponding p-value is at most 0.05. A similar figure based on Pearson correlations is depicted in Figure 3.

### 6 Strand dependence of vIP values

| Triplets | Reference strand | | | Strand of minimum vIP | | |
|---|---|---|---|---|---|---|
| | $S_{SC}$ | $S_{SN}$ | $S_G$ | $S_{SC}$ | $S_{SN}$ | $S_G$ |
| silent | -0.63 | -0.64 | -0.63 | -0.74 | -0.76 | -0.73 |
| missense | -0.56 | -0.56 | -0.53 | -0.67 | -0.65 | -0.59 |
| nonsense | -0.51 | -0.46 | -0.45 | -0.63 | -0.55 | -0.51 |
| intronic | -0.50 | -0.52 | -0.54 | -0.55 | -0.60 | -0.65 |
| UTR | -0.56 | -0.55 | -0.55 | -0.65 | -0.65 | -0.62 |
| splice region | -0.43 | -0.50 | -0.47 | -0.53 | -0.63 | -0.56 |
| intergenic | -0.53 | -0.53 | -0.60 | -0.61 | -0.61 | -0.69 |

(a)

| Quintuplets | Reference strand | | | Strand of minimum vIP | | |
|---|---|---|---|---|---|---|
| | $S_{SC}$ | $S_{SN}$ | $S_G$ | $S_{SC}$ | $S_{SN}$ | $S_G$ |
| silent | -0.60 | -0.61 | -0.59 | -0.66 | -0.68 | -0.62 |
| missense | -0.56 | -0.58 | -0.55 | -0.62 | -0.62 | -0.56 |
| nonsense | -0.21 | -0.40 | -0.23 | -0.25 | -0.45 | -0.27 |
| intronic | -0.46 | -0.49 | -0.54 | -0.50 | -0.54 | -0.63 |
| UTR | -0.55 | -0.53 | -0.57 | -0.59 | -0.60 | -0.60 |
| splice region | -0.35 | -0.41 | -0.34 | -0.45 | -0.50 | -0.42 |
| intergenic | -0.51 | -0.47 | -0.61 | -0.56 | -0.52 | -0.64 |

(b)

Table S5: Pearson correlation coefficients $r$ between $F$, the logarithm of the normalized frequency of each mutated motif, and the vIP of (a) triplets and (b) quintuplets in the three datasets $S_{SC}$, $S_{SN}$, $S_G$. In the columns called "Reference strand", $F$ and vIP values were computed on the DNA strand annotated in the dataset; in the columns "Strand of minimum vIP", they are computed on the strand of minimal vIP. These results must be compared with those in Table 2.a-b of the main manuscript, where $F$ and vIP values are averaged over the two DNA strands.

#### 7 Spearman correlations for somatic and germline mutations

| Triplets - NXN | | | | |
|---|---|---|---|---|
| | $S_{SC}$ | $S_{SN}$ | $S_G$ | Total |
| silent | -0.82 | -0.85 | -0.87 | -0.84 |
| missence | -0.72 | -0.73 | -0.63 | -0.72 |
| nonsence | -0.66 | -0.57 | -0.57 | -0.68 |
| intronic | -0.57 | -0.64 | -0.66 | -0.60 |
| UTR | -0.68 | -0.66 | -0.69 | -0.67 |
| splice_region | -0.54 | -0.55 | -0.52 | -0.56 |
| intergenic | -0.63 | -0.63 | -0.75 | -0.64 |

(a)

| Quintuplets - NNXNN | | | | |
|---|---|---|---|---|
| | $S_{SC}$ | $S_{SN}$ | $S_G$ | Total |
| silent | -0.74 | -0.78 | -0.74 | -0.75 |
| missense | -0.68 | -0.71 | -0.65 | -0.68 |
| nonsense | -0.29 | $-0.44^{\times}$ | -0.32 | -0.33 |
| intronic | -0.43 | -0.51 | -0.62 | -0.48 |
| UTR | -0.62 | -0.60 | -0.68 | -0.64 |
| splice region | -0.41 | $-0.47^{\times}$ | -0.37 | -0.40 |
| intergenic | -0.55 | -0.48 | $-0.75^{\times}$ | -0.50 |

(b)

Table S6: Spearman correlation coefficients $\rho$ between $\overline{F}$ and $\overline{\text{vIP}}$ values of (a) triplets and (b) quintuplets in the three datasets $S_{SC}$, $S_{SN}$, $S_G$ containing somatic cancer mutations, somatic non-disease mutations and germline mutations, respectively. "Total" means the ensemble of the three datasets. The p-values are all lower than 0.001. A $\times$ superscript indicates that the dataset contains less than 50 times the number of triplets (which is 64; in (a)) or of quintuplets (1024; in (b)); the dataset content is specified in Supplementary Table 2. We disregard these values with low statistical significance in our analyses. Similar Tables with Pearson correlations are found in Table 2 of the main manuscript.

### 8 Comparison with previous analyses

In previous papers [5, 6], we already computed the correlation between vIP and the normalized mutation frequencies of the $S_G$ set of germline mutations reported in ClinVar [7], and found slightly different results with, in general, slightly weaker anticorrelations. The reasons of these differences are described below:

- An earlier version of the ClinVar dataset was used;
- Only silent mutations were considered in [6];
- The Pearson correlation computed in [5] was between vIP values and mutational motif frequencies rather than between vIP values and the logarithm of mutational motif frequencies.
- The normalization of mutational motif frequencies was performed based on the frequency of the motifs in the genome regions in which they occur (exon, intron, UTR) in [5, 6]. Here the normalization was performed independently of the type of genome region, as the electron hole migration along the DNA stack is blind to it.
- The mutational motif frequencies were considered on the DNA strand annotated in the dataset in [5, 6], and thus the correlations were performed between vIP values and mutational motif frequencies or their logarithm rather than between the averages over both strands, as done here (see Section 9 herabove).

The rationale behind choosing to compute $\overline{\text{vIP}} - \overline{F}$ correlations rather than $\text{vIP} - F$ is that it is impossible to determine on what DNA strand the mutation primarily occurred, and what strand was mutated to restore a correct base pairing.

### 9 $\overline{\text{vIP}} - \overline{F}$ correlations per chromosome

| chromosome | $r(\overline{F}, \overline{\text{vIP}})$ | N |
|---|---|---|
| chr1 | -0.78 | 183,844 |
| chr2 | -0.76 | 151,935 |
| chr3 | -0.76 | 102,590 |
| chr4 | -0.75 | 68,353 |
| chr5 | -0.76 | 90,875 |
| chr6 | -0.77 | 88,040 |
| chr7 | -0.77 | 92,166 |
| chr8 | -0.77 | 66,766 |
| chr9 | -0.78 | 75,722 |
| chr10 | -0.78 | 68,794 |
| chr11 | -0.77 | 114,769 |
| chr12 | -0.78 | 90,436 |
| chr13 | -0.73 | 32,897 |
| chr14 | -0.77 | 57,929 |
| chr15 | -0.77 | 59,925 |
| chr16 | -0.79 | 85,765 |
| chr17 | -0.78 | 103,324 |
| chr18 | -0.75 | 31,396 |
| chr19 | -0.78 | 123,420 |
| chr20 | -0.79 | 47,526 |
| chr21 | -0.77 | 20,545 |
| chr22 | -0.79 | 38,835 |
| chrX | -0.77 | 69,896 |
| chrY | -0.64 | 446 |

Table S7: Pearson correlation $r$ between $\overline{\text{vIP}}$ and $\overline{F}$ per chromosome for silent mutations in the three datasets $S_{SC}$, $S_{SN}$, $S_G$ taken together; N is the number of mutations. Here, the motif frequency $\hat{f}$ appearing in Eq. (1) of the main text is computed for each chromosome separately.

### 10 $\overline{\text{vIP}} - \overline{F}$ correlations according to the tissue

| Disease | Triplet | Quintuplet |
|---|---|---|
| lung | -0.87 | -0.79 |
| kidney | -0.87 | |
| liver | -0.86 | -0.76 |
| ovary | -0.85 | |
| thyroid | -0.85 | |
| small_intestine | -0.85 | |
| soft_tissue | -0.85 | |
| bone | -0.84 | |
| upper_aerodigestive_tract | -0.84 | $-0.75^{\times}$ |
| biliary_tract | -0.83 | |
| haematopoietic_and_lymphoid_tissue | -0.83 | $-0.76^{\times}$ |
| prostate | -0.82 | |
| pancreas | -0.82 | |
| large_intestine | -0.81 | -0.74 |
| oesophagus | -0.81 | $-0.72^{\times}$ |
| central_nervous_system | -0.81 | $-0.73^{\times}$ |
| endometrium | -0.80 | -0.69 |
| stomach | -0.80 | -0.74 |
| breast | -0.79 | $-0.71^{\times}$ |
| urinary_tract | -0.78 | -0.70 |
| skin | -0.76 | -0.69 |
| cervix | -0.76 | |
| nervous system | -0.72 | |

Table S8: Pearson correlation $r$ between $\overline{\text{vIP}}$ and $\overline{F}$ for silent mutations in the set $S_{SN}$ of cancer mutations, according to the tissues were they are found as annotated in COSMIC [2]. Tissues were sorted by decreasing correlation values for the triplet motifs. A $\times$ superscript indicates that the dataset contains between 25 and 50 times the number of triplets (which is 64) or of quintuplets (which is 1024); correlations based on less than 25 times the number of motifs are not reported.
